# Separation of newly replicated bacterial chromosomes: the role of *Escherichia coli* Topoisomerase IV

**DOI:** 10.1101/2020.02.14.948935

**Authors:** Emily Helgesen, Frank Sætre, Kirsten Skarstad

## Abstract

Topoisomerase IV (TopoIV) is a vital bacterial enzyme which disentangles newly replicated DNA and enables segregation of daughter chromosomes. In bacteria, DNA replication and segregation are concurrent processes. This means that TopoIV must continually remove inter-DNA linkages during replication. There exists a short time lag of about 5-10 minutes between replication and segregation in which the daughter chromosomes are intertwined. Exactly where TopoIV binds during the cell cycle has been the subject of much debate. We show here that TopoIV localizes to the origin proximal side of the fork trailing protein SeqA and follows the movement pattern of the replication machinery in the cell.

## Introduction

Proper segregation of newly replicated DNA is essential for the viability and genetic stability of all cell types. Due to the superhelical nature of DNA molecules, topology challenges are inevitable during the process of DNA replication, as the template strands are separated and duplicated. More specifically, tension arises in front of the replication machinery (hereafter called the replication fork) as the parental DNA strands are pulled apart, which results in the formation of positive supercoils (overwinding) ^1, 2^. Some of these positive supercoils may diffuse towards the newly replicated DNA molecules behind the replication fork, and the replication fork most likely rotates to alleviate some of the topology tension piling up ahead ^3^. As a consequence, the newly replicated DNA molecules become intertwined, and this type of entanglement is typically referred to as precatenanes ^1-4^. Without the removal of precatenane linkages it becomes impossible for the cell to segregate the DNA prior to cell division. Highly specific mechanisms therefore exist to resolve the topological issues that arise during DNA replication, and at the core of these mechanisms we find the enzymes categorized as type II topoisomerases ^2^.

In *Escherichia coli* two type II topoisomerases are involved in enabling both DNA replication and timely DNA segregation, namely Gyrase and Topoisomerase IV (TopoIV). Both of these enzymes work by first performing a transient double strand break in one molecule, then leading a second DNA duplex through the cut and lastly, resealing the cut. They are heterotetrameric structures consisting of GyrA and GyrB subunits or ParC and ParE subunits for Gyrase and TopoIV, respectively. The GyrA/ParC subunit contains the DNA binding and catalytical properties of the enzyme, whereas ATP binding resides in GyrB/ParE ^5^. It is now generally well recognized that Gyrase acts in front of the replication fork to remove excess positive supercoiling to support fork progression, whereas TopoIV mainly removes precatenane linkages after replication to facilitate DNA segregation ^6-9^. However, there has been much debate concerning the precise timing and localization of TopoIV action. It has been suggested that TopoIV activity is limited to the D-period (when a round of DNA replication is completed, see supplementary figure 1) and that TopoIV localizes mainly at the terminus ^10^. It has also been indicated that the catalytically active TopoIV molecules bind in clusters at the origins, where they are recruited and stimulated by MukB, an SMC (structural maintenance of chromosomes) protein ^11-13^. Moreover, there is a time lag of 5-10 minutes between replication of the DNA and segregation of the newly replicated DNA, which is termed the “cohesion period” ^6, 8, 9, 14^. Whether this means that TopoIV does not immediately gain access to the DNA after replication (i.e. that precatenanes hold the homologous DNA together), or if other factors such as proteins bridging the DNA is causing this delay, is not completely understood.

**Figure 1:**
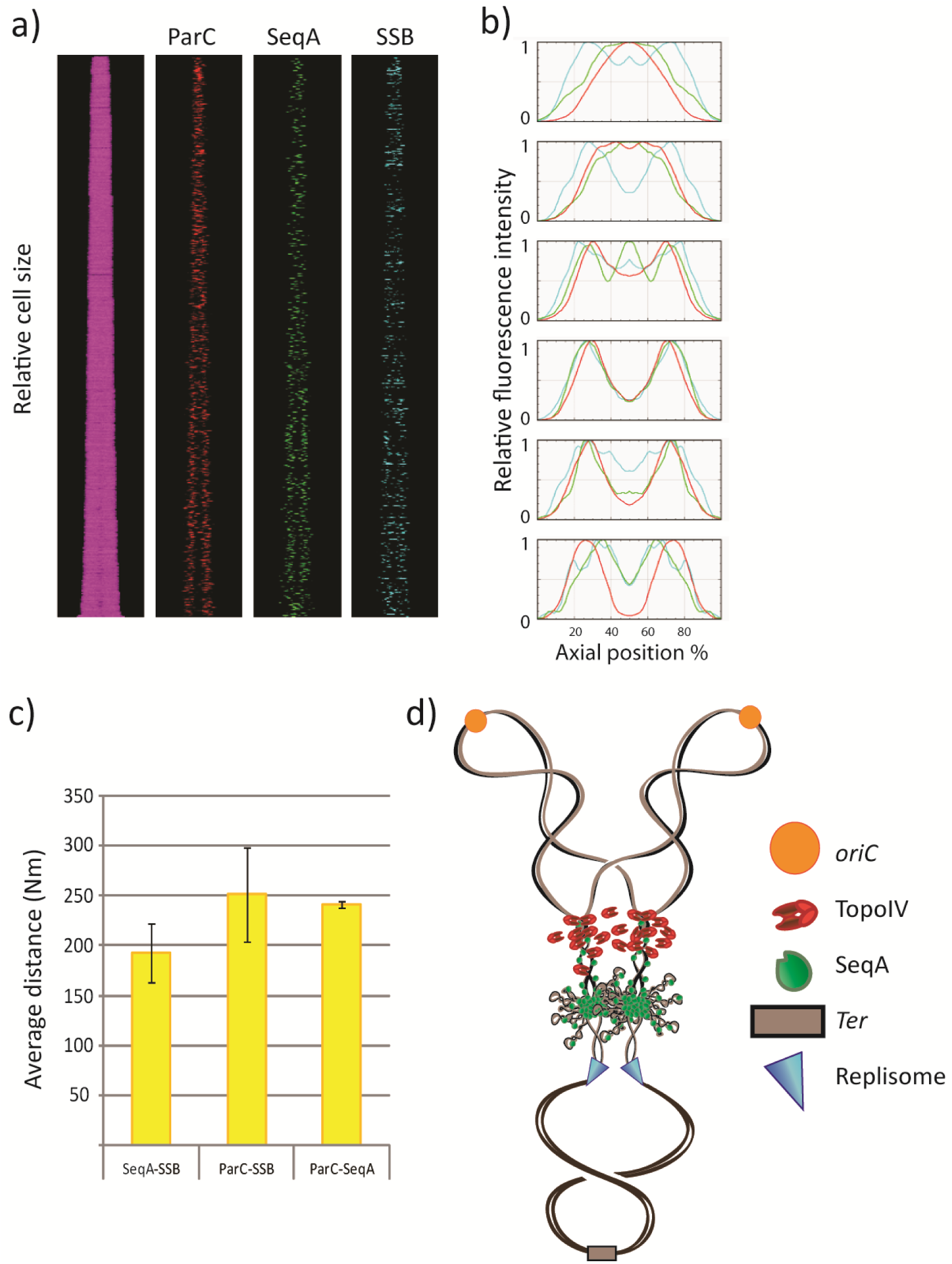
Fluorescence microscopy indicates that TopoIV trails the replication forks and primarily decatenates the precatenanes farthest from the replisome. a) Kymographs showing fluorescent foci of TopoIV (ParC-mKate2), SeqA (SeqA-YFP) and replisome (SSB-CFP) in cells stacked horizontally according to cell size (smallest cells top and largest cells bottom). b) Plots of relative fluorescence intensity (Y-axis) according to position on the cell long axis (X-axis) in groups of cell sizes/ages from smallest (top) to largest (bottom). b) Plot showing the average distances (nm) between SeqA-SSB, ParC-SSB and ParC-SeqA in the cells from three independent experiments. Error bars are included in the plot. d) Hypothetical cartoon showing the relative localization of the replisomes (cyan triangles), SeqA molecules (green) and TopoIV molecules (red) on an actively replicating chromosome. The old DNA is shown as black lines, the newly synthesized DNA as grey lines, the origins (*oriC*s) in orange and the terminus region in brown.

In this work we have sought to elucidate the localization and movement of TopoIV with respect to the replication fork and a fork-trailing protein named SeqA. SeqA is a negative modulator of replication initiation, which binds to newly replicated, hemimethylated GATC-sites ^15-17^. SeqA forms multimeric structures which trail the replication forks dynamically, always binding to the newest DNA ^18-21^. The SeqA-DNA complexes are large and typically encompass 100 kb of DNA. We have previously found that the SeqA multimer binds at a distance from the replisome (on average 200-300 nm) ^22^. The newly replicated DNA molecules were found to be kept close together on this stretch, i.e. they were cohesed. The localization of the cohesed DNA and the replisomes in the cell were visualized by utilizing fluorescently tagged SeqA (SeqA-YFP) and replisome proteins (SSB-CFP), respectively.

We find here that fluorescently tagged TopoIV (ParC-mKate2) exhibits a localization pattern throughout the cell cycle compatible with the model that TopoIV trails SeqA and the replisome during replication. Moreover, the average distance between TopoIV and the replisome is always larger than that between SeqA and the replisome. This indicates that TopoIV is indeed excluded from binding to the DNA immediately after its replication. Inhibition of TopoIV using a fluoroquinolone antibiotic, Ciprofloxacin, lead to an increased distance between SeqA and TopoIV, presumably because the TopoIV molecules become “stuck” in DNA ternary complexes, thereby lagging even further behind the replication machinery.

## Results and discussion

### TopoIV most likely trails SeqA during DNA replication

In order to investigate the localization of TopoIV with respect to the replication fork and the newly replicated DNA, we constructed a strain which contains fluorescent tags on the single-stranded binding protein (SSB-CFP) present at the replisome, on SeqA (SeqA-YFP) and on TopoIV (ParC-mKate2). The cells exhibited a normal growth rate and cell cycle compared to the wild type background, i.e. they were able to successfully complete DNA replication and had no observable segregation issues (see Table 1 for generation times and cell cycle parameters and Fig S1 for flow cytometry histograms). We grew the cells in poor medium (acetate medium) to early exponential phase (OD∼0.15) and investigated the living cells with snap-shot fluorescence microscopy. The images were subjected to analysis with Coli Inspector (see Methods for details) in order to assess the positioning of fluorescent foci. From kymographs of the fluorescent foci (in which the cells are stacked according to cell size) (Fig 1a) and plots of relative fluorescence intensity according to position along the cell long-axis (Fig 1b), we found that TopoIV had a localization pattern that resembled that of SeqA and the replisome. This supports the model that TopoIV trails the replication machinery to ensure processive removal of precatenanes, and that it is not restricted to performing decatenation after replication termination. Flow cytometry analysis of DNA content (as described in ^23^) showed that the cells had a cell cycle in which the newborn cell contained one replicating chromosome where the replication forks were about to terminate (see Fig S1 for DNA histograms and schematic cell cycle cartoons). Cells which are about to terminate replication of a chromosome have already segregated their two origins to the respective quarter positions in the cell ^22, 24^. In this study we find that TopoIV is localized at mid-cell at this stage of the cell cycle, i.e. in the newborn cells (Fig 1b top panel). This indicates that TopoIV is not exclusively found in clusters associated with MukB at the origins, as inferred in ^11, 13^. Recently, it was found that the MukB-TopoIV interaction in fact promoted DNA condensation and did not involve any catalytic activity of TopoIV ^25^. It may therefore be that TopoIV bound to MukB at origins does not contribute to resolution of precatenanes. The reason for the discrepancy between our study and previous studies of TopoIV localization is not known.

**Table 1.** Cell cycle parameters for strains grown in acetate medium at 28oC (averages from at least three separate experiments +/- SEM)

| Strain | Doubling time | C-period | D-period | Initiation |
| --- | --- | --- | --- | --- |
| **AB1157**<br>Parent | 194 +/-26 | 103 +/-11 | 122 +/-6 | 164 +/-47 |
| **EH29**<br>SeqA-YFP<br>SSB-CFP<br>ParC-mKate2 | 184 +/-12 | 104 +/-10 | 126 +/-7 | 139 +/-25 |
| **EH32**<br>gyrA<sup>L83Y87</sup> | 177 +/-42 | 119 +/-54 | 73 +/-44 | 93 +/-92 |
| **EH34**<br>SeqA-YFP<br>ParC-mKate2<br>gyrA<sup>L83Y87</sup> | 156 +/-36 | 96 +/-23 | 86 +/- 11 | 68 +/-58 |

As observed in previous studies the replisome appears to be more dynamic compared to the replisome-trailing SeqA structures, as one replisome focus more frequently represents one replication fork at each of the quarter positions in cells growing with one replicating chromosome ^22, 26^ (see young cells in Fig 1a and b) compared to SeqA which stays at mid cell (thus representing four strands of newly replicated DNA). We found that TopoIV had a localization pattern which was more similar to that of SeqA than to that of the replisome, which is especially prominent in the newborn cells harboring SeqA and TopoIV at midcell (Fig 1b, top panels). This may indicate that TopoIV is closer to SeqA than to the replisome. To further elucidate this scenario we measured the distances between the three fluorescently tagged structures using high-throughput image analysis scripts described previously ^22, 24^. Briefly, after processing of the images, the script measures the distances between the highest intensity pixels from each channel/focus, in which the highest concentration of molecules are likely to be situated. From three separate experiments we found that the average distance between SeqA and the replisome was always less than that between TopoIV and the replisome (see average values Fig 1c). This finding suggests that TopoIV binds on the origin-proximal side of SeqA and the replisome (Fig 1d).

### The distance between SeqA and TopoIV increases when TopoIV is inhibited by Ciprofloxacin

The group of antibiotics termed fluoroquinolones is known to bind and inhibit Gyrase and TopoIV by forming a ternary complex with these enzymes and DNA. Upon drug interaction Gyrase/TopoIV remains as a “frozen” adduct on DNA after the cleavage step, and is unable to reseal the double-strand ends after strand passage ^27^. We decided to use the fluoroquinolone Ciprofloxacin to shed more light on the positioning of TopoIV during replication. If TopoIV is localized between the SeqA complex and the replisome, one would expect to observe a perturbation of SeqA focus formation upon inhibition of TopoIV, since SeqA may “collide” into the frozen adducts that occupy the space necessary for SeqA binding and multimerization. If, on the other hand, TopoIV trails SeqA, it would be expected that the distance between the SeqA complex and TopoIV increases compared to the untreated control, as the TopoIV-Ciprofloxacin adducts will be lagging behind on the origin-proximal side of the SeqA complex.

To ensure that only TopoIV would be targeted in our experiments, we used a strain which contains two mutations in the GyrA subunit of Gyrase (L83 and Y87) ^28^, rendering Gyrase insensitive to fluoroquinolones, in addition to the fluorescently tagged SeqA (SeqA-YFP) and TopoIV (ParC-mKate2) constructs. The cells were grown in acetate medium to early exponential phase (OD∼0.15) and either imaged directly (as described in the previous section) or treated with 0.1 µg/ml Ciprofloxacin for 45 minutes prior to imaging.

Image analysis showed that the localization pattern of SeqA and TopoIV was different in the Ciprofloxacin-treated cells compared to the untreated control (Fig 2a). This is not surprising, considering that the ability of TopoIV to properly facilitate decatenation and segregation is compromised. However, the Ciprofloxacin-treated cells had no problem with SeqA focus formation, and when measuring the SeqA-TopoIV distances in the cells we found that the average distance was indeed increased in the Ciprofloxacin-treated culture (r= .70, p= .033) (Fig 2b). A schematic model is depicted in Fig 2c, showing how the Ciprofloxacin-bound TopoIV complexes may become stuck in the DNA and lag behind SeqA, thus leading to an increased SeqA-TopoIV distance. The result supports our previous inferences and strengthens the theory that TopoIV is excluded from the DNA “cohesion window” between SeqA and the replisome.

**Figure 2:**
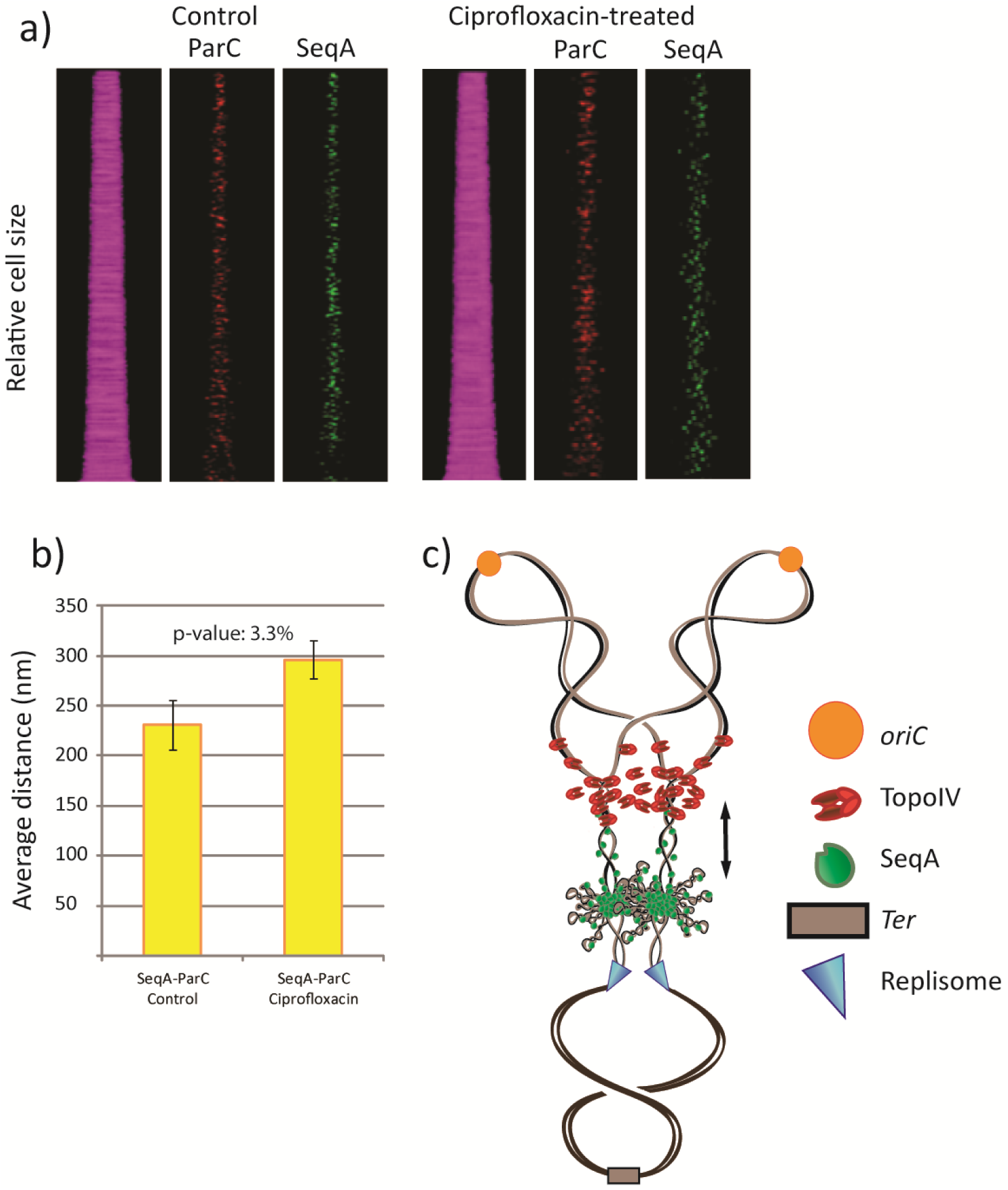
Treatment with the antibiotic Ciprofloxacin supports the idea that TopoIV activity trails the replication fork on the origin-proximal side of the «cohesion window». a) Kymographs showing fluorescent foci of TopoIV (ParC-mKate2) and SeqA (SeqA-YFP) in cells stacked horizontally according to cell size (smallest cells top and largest cells bottom). Untreated cells are shown to the left whereas cells treated with Ciprofloxacin (0.1 µg/ml) for 45 minutes are shown to the right. b) Plot showing the average distances (nm) between SeqA and TopoIV (ParC) in untreated (left) and Ciprofloxacin treated (right) cells from three independent experiments. Error bars are included in the plot. The p-value for increase in SeqA-ParC distances in Ciprofloxacin treated cells is indicated in the plot and was calculated using a paired, one-tailed T-test on average distances from three independent experiments. c) Hypothetical cartoon showing how TopoIV (red) may lag farther behind SeqA (green) when inhibited by Ciprofloxacin on the newly replicated DNA. The old DNA is shown as black lines, the newly synthesized DNA as grey lines, the origins (*oriC*s) in orange and the terminus region in brown.

How this cohesion window is maintained is currently not understood. One possible explanation could be that this stretch of DNA is occupied by other proteins, which inhibit TopoIV binding. For instance, high concentrations of SeqA have been shown to inhibit TopoIV activity ^29^, and it has been suggested that SeqA clusters protect the intercatenation linkages from TopoIV ^6^. Moreover, an interaction between SeqA and TopoIV has been indicated, and lower concentrations of SeqA seem to stimulate TopoIV activity ^29^. It could therefore be that TopoIV is directed to a DNA region with fewer molecules of SeqA, as indicated in Fig 1d. SeqA plays an important role in regulating the hemimethylation status of newly replicated DNA and it is striking that the period of hemimethylation is similar to that of the cohesion period ^14, 15, 30^. It could also be that the topology of the precatenated DNA directly behind the replication fork is suboptimal for TopoIV binding.

Speculation set aside; there are certainly clear advantages of keeping the newly replicated sister DNA close together for a period of time. It allows for various vital processes to occur due to the proximity of the two homologous double-strands, such as homologous recombination, replication fork remodeling, reversal and restart reactions ^8, 22, 31^.

## Methods

Strain construction: All strains used in experiments are derivatives of the *E. coli* K-12 strain AB1157 ^32^ and are listed in supplementary Table S1. Localization studies of SeqA were done with cells containing the yellow fluorescent protein (YFP) fused to the C-terminal end of SeqA ^33^. The *seqA-yfp* gene was expressed from the endogenous chromosomal promoter. The YFP protein was from ^34^ and connected to SeqA via a four-amino acid linker ^18^. Studies of SSB localization were with cells containing the SSB-CFP allele inserted in place of the *E. coli lamB* gene and was kindly provided by A. Wright (G. Leung et al., unpublished) ^22^. The cells contained the wild-type *ssb* gene on the chromosome. The fluorescent version of ParC was constructed in this study. Briefly, the *mKate2* gene was PCR amplified from the plasmid pTEC20 ^35^, subcloned via pGEM-T easy (Promega) and inserted upstream of a chloramphenicol resistance cassette in the plasmid pSF36 (pUC19+cm-FRT-*HindIII*) yielding plasmid pEH04. Primers with 50 bp homology to the C-terminus of *parC* and 50 bp homology to the sequence directly downstream of the *parC* gene were used to amplify *mKate2* with *parC* homology tails from pEH04. These were as follows:

5’GTGTTGAGATCGACTCTCCTCGCCGTGCCAGCAGCGGTGATAGCGAAGAG TCTGGTTCTGGTTCTGGTTCTGGTTCTGGTTCTGGT GTGAGCGAGCTGATTAAGGAG 3’ 5’TCATCCGGCGTTCCTTGCAAGCGGGAGGAAACAGCGCCCTCCCCGGCATATTACGCCAAGCTTGTGTAGGCT 3’

Next, the PCR-product with flanking tails was electroporated into AB1157 cells, and homologous recombination facilitated by induction of plasmid pRed/ET (GeneBridges) as described in ^36^.

The construct was verified by sequencing to be inserted at the correct position on the chromosome (at the endogenous *parC* gene) and to contain an amino acid linker sequence (Ser-Gly)^6^ between the C-terminal of ParC and the start of mKate2. To obtain the strains used here with combinations of fluorescent constructs and/or mutations, P1 transduction ^37^ and FLP recombinase (pCP20) was used ^38^.

Cell growth: Cells were grown at 28°C in AB minimal medium ^39^ supplemented with 0.4% sodium acetate, 1-μg ml^−1^ thiamine, 80-μg ml^−1^ threonine, 20-μg ml^−1^ leucine, 30-μg ml^−1^proline, 22-μg ml^−1^ histidine and 22-μg ml^−1^ arginine (acetate medium). The doubling time (τ) was found by optical density (OD) measurements. Cells were grown to OD ∼ 0.15 (early exponential phase) at which time they were prepared for flow cytometry analysis or fluorescence microscopy. For experiments with Ciprofloxacin, EH34 cells were treated with 0.1 ug/ml for 45 minutes prior to imaging.

Flow cytometry and cell cycle analysis: Exponentially growing cells were fixed in ethanol or treated with 300-μg/m rifampicin and 10-μg/ml cephalexin to inhibit replication initiation ^40^ and cell division ^41^, respectively. Growth of drug-treated samples continued for 3–4 generations, after which they were fixed in ethanol. Drug-treated cells ended up with an integral number of chromosomes ^40^, which represents the number of origins at the time of drug treatment (replication run-out). Flow cytometry was performed as previously described ^42^ using an LSR II flow cytometer (BD Biosciences) and FlowJo 7.2.5 software. Cell cycle parameters, numbers of origins and replication forks per cell were obtained by analysis of the DNA distributions obtained by flow cytometry as described ^23^.

Fluorescence microscopy imaging: for fluorescence microscopy exponentially growing cells were immobilized on an agarose pad (1% agarose in phosphate-buffered saline) and covered with a #1.5 coverslip. Images were acquired with a Leica DM6000 microscope equipped with a Leica EL6000 metal halide lamp and a Leica DFC350 FX monochrome CCD camera. Phase contrast imaging was performed with an HCX PLAPO 100x/1.40 NA objective. Narrow band-pass filter sets (CFP: Ex BP436/20, Em BP480/40, YFP: Ex BP510/20, Em BP560/40, Cy3: Ex BP545/30, Em BP610/75) were used for fluorescence imaging.

During image acquisition, saturated pixels were avoided. The raw images were saved for further image processing (see below).

Image processing and analysis: imaging adjustments (brightness and contrast) were performed in Image J or Fiji software. We used the public domain Coli-Inspector project to obtain fluorescence intensity profiles of the cells and to do vertical plotting of fluorescence and phase contrast images of cells. Coli-Inspector runs under ImageJ/Fiji in combination with the plugin ObjectJ (http://simon.bio.uva.nl/objectj/). The average fluorescence intensity profile of cells was plotted against the cell long axis, in groups of increasing cell length, as described ^43^. Vertical plotting of cells was done in the order of gradual increase in cell length. Age classes of cells were defined by the cell length, assuming that length increases linearly.

We used a Python-based script developed in our group for measurements of distances between neighboring spots/foci that are registered in two different fluorescence channels. The script outputs all registered distances (in this case distances between SeqA, ParC and SSB) per cell, and these values were used to calculate average distances from at least three separate experiments. Image processing for automated analysis using this script was performed in Image J using the following tools: i) Background subtraction with default Rolling disk (diameter 10 pixels), ii) Deconvolution using the Richardson-Lucy algorithm (100 iterations), iii) Median filter, iv) thresholding by Max Entropy (see ^22^ for details). The positive correlation and p-value for increase in SeqA-ParC distances in Ciprofloxacin treated cells was calculated using a paired, one-tailed T-test on average distances from three independent experiments.

## Acknowledgements

We thank A. Wahl for excellent technical assistance, B. Sen for construction of strains and S. Fossum-Raunehaug for the plasmid pSF36. The Flow Cytometry Core Facility (by T. Stokke) at The Norwegian Radium Hospital is greatly acknowledged for help with flow cytometry analysis. We thank A. Wright, M. Radman and L. Zechiedrich for providing strains. This work was supported by a grant from Helse Sør-Øst (project 2012062).

## Author contributions

Conceived and developed the study: E.H., F.S., K.S. Performed the experiments and analyzed the data: E.H., F.S. Interpreted the results: E.H., K.S. Wrote the manuscript: E.H., K.S.

