## Supplementary material for "Separation of newly replicated bacterial chromosomes: the role of *Escherichia coli* Topoisomerase IV": Table S1 and Fig S1

**Table S1: Table of strain names, relevant features and sources.**

| **Strain name** | **Relevant features**[a] | **Source** |
|---|---|---|
| AB1157 | Wild type strain | (1) |
| GL224 | MG1655 *SSB-CFP::cat* in *lamB* | A. Wright and G. Leung |
| MG1655 *seqA-YFP* | MG1655 *seqA-YFP::cat* | (2) |
| BS30 | AB1157 *parC-mKate2::cat* | This work |
| BS36 | AB1157 *seqA-YFP parC-mKate2::cat* | This work |
| EH29 | AB1157 *seqA-YFP parC-mKate2 SSB-CFP::cat* | This work |
| LZ3099 | C600 *gyrA*$^{S83L+D87Y}$ | (3) |
| EH32 | AB1157 *gyrA*$^{S83L+D87Y}$ | This work |
| EH34 | AB1157 *seqA-YFP parC-mKate2::cat gyrA*$^{S83L+D87Y}$ | This work |

[a] Description of strain constructions in Materials and Methods

**Figure S1: Flow cytometry histograms and schematic cell cycle cartoon for EH29 cells**

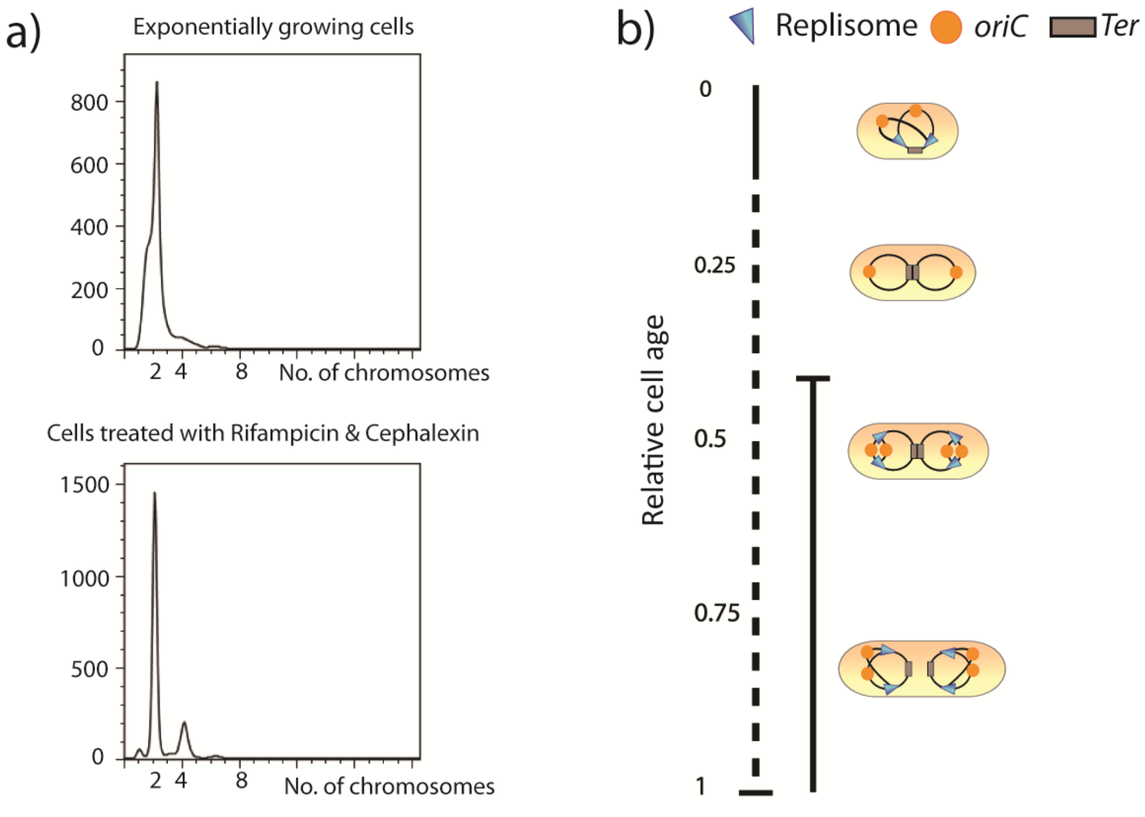


Fig S1: a) Flow cytometry histograms showing the DNA content of exponentially growing cells (top panel) and cells treated with Rifampicin and Cephalexin to produce a run-out of

DNA replication (bottom panel) of the strain EH29. b) Schematic cartoon showing cell cycle progression and DNA content according to cell size. The solid black line represents the replication period (C-period) with initiation around a relative cell age of 0.42, whereas the stippled black line represents the post replication period (D-period). Cell cartoons are shown to indicate DNA content and cell cycle progression. The cells were grown at 28$^{o}$C in acetate medium (see Methods for medium composition).


1. Howard-Flanders, P., Simson, E. and Theriot, L. (1964) A locus that controls filament formation and sensitivity to radiation in *Escherichia coli* K-12. *Genetics*, **49**, 237-246.
2. Babic, A., Lindner, A.B., Vulic, M., Stewart, E.J. and Radman, M. (2008) Direct visualization of horizontal gene transfer. *Science (New York, N.Y.)*, **319**, 1533-1536.
3. Morgan-Linnell, S.K. and Zechiedrich, L. (2007) Contributions of the combined effects of topoisomerase mutations toward fluoroquinolone resistance in *Escherichia coli*. *Antimicrobial agents and chemotherapy*, **51**, 4205-4208.
